# De novo virus inference and host prediction from metagenome using CRISPR spacers

**DOI:** 10.1101/2020.09.04.282665

**Authors:** Ryota Sugimoto, Luca Nishimura, Phuong Nguyen Thanh, Jumpei Ito, Nicholas F. Parrish, Hiroshi Mori, Ken Kurokawa, Hirofumi Nakaoka, Ituro Inoue

**Affiliations:** Human Genetics Laboratory, National Institute of Genetics, Research Organization of Information and Systems, 1111 Yata, Mishima, Shizuoka, Japan; The Graduate University for Advanced Studies, SOKENDAI, 1111 Yata, Mishima, Shizuoka, Japan; Division of Systems Virology, Department of Infectious Disease Control, International Research Center for Infectious Diseases, Institute of Medical Science, The University of Tokyo, 4-6-1 Shirokanedai, Minato-ku, Tokyo, Japan; Genome Immunobiology RIKEN Hakubi Research Team, Center for Integrative Medical Sciences, RIKEN, 1-7-22 Suehiro-cho, Tsurumi-ku, Yokohama, Kanagawa, Japan; Genome Evolution Laboratory, National Institute of Genetics, Research Organization of Information and Systems, 1111 Yata, Mishima, Shizuoka, Japan; Department of Cancer Genome Research, Sasaki Institute, 2-2 Kandasurugadai, Chiyoda-ku, Tokyo, Japan

## Abstract

Viruses are the most numerous biological entity, existing in all environments and infecting all cellular organisms. Compared with cellular life, the evolution and origin of viruses are poorly understood; viruses are enormously diverse and most lack sequence similarity to cellular genes. To uncover viral sequences without relying on either reference viral sequences from databases or marker genes known to characterize specific viral taxa, we developed an analysis pipeline for virus inference based on clustered regularly interspaced short palindromic repeats (CRISPR). CRISPR is a prokaryotic nucleic acid restriction system that stores memory of previous exposure. Our protocol can infer viral sequences targeted by CRISPR and predict their hosts using unassembled short-read metagenomic sequencing data. Analysing human gut metagenomic data, we extracted 11,391 terminally redundant CRISPR-targeted sequences which are likely complete circular genomes of viruses or plasmids. The sequences include 257 complete crAssphage family genomes, 11 genomes larger than 200 kilobases, 766 genomes of *Microviridae* species, 114 genomes of *Inoviridae* species and many entirely novel genomes of unknown taxa. We predicted the host(s) of approximately 70% of discovered genomes by linking protospacers to taxonomically assigned CRISPR direct repeats. These results support that our protocol is efficient for de novo inference of viral genomes and host prediction. In addition, we investigated the origin of the diversity-generating retroelement (DGR) locus of the crAssphage family. Phylogenetic analysis and gene locus comparisons indicate that DGR is orthologous in human gut crAssphages and shares a common ancestor with baboon-derived crAssphage; however, the locus has likely been lost in multiple lineages recently.

## Introduction

Viruses are not included among the domains of life, but they are the most numerous biological entity, presumably containing the most genetic material on Earth. The number of viruses infecting microbial populations is estimated at 10^31^1^. Viruses exist in all environments and infect all cellular organisms. In the historical experiments of Hershey and Chase^2^, a virus was used to first show that DNA is the genetic material, and the first sequenced complete genome was viral^3,4^; despite those historical efforts to obtain genetic information, the evolution and origins of viruses are poorly understood, reasons for which are related to the nature of viruses. Firstly, viral genomes are extremely diverse in size, ranging from several thousand to a few million bases, and in genomic content, encode various non-overlapping sets of genes. In fact, there are no genes universally present among all viruses, indicating that a monophyletic explanation of viral origin is implausible^5,6^. Secondly, most viral genes show little to no homology to cellular genes. Despite their parasitic nature, viral genomes contain sparse sequences from which to infer shared lineage or evolutional history with cellular life. Considering these confounding biological features, we were motivated to uncover viral genomic sequences without relying on either marker genes or known sequences.

Metagenomics^7^ became a popular method to sequence viral genomes. Metagenomics sequences all genetic materials extracted from a given organism or tissue. The advantage of a metagenomic approach is, with sufficient depth of sequencing, presumably all genetic information of viruses in the sample can be obtained. However, because metagenomic approaches obtain a mixture of cellular and viral genomic information, it is necessary to subsequently distinguish viral from cellular sequences.

Several methods have been developed to detect viral sequences from metagenomic data, after assembling sequences into those derived from presumably contiguous DNA molecules called contigs. One virus detection software, VirSorter^8^ detects regions within contigs enriched with viral ‘hallmark genes’, viral-like genes, uncharacterized genes, and shortened genes. To detect viral proteins, this program uses a custom-made protein database including both reference viral proteins and predicted proteins from virome samples. Another virus detection method, VirFinder^9^ is a k-mer based machine learning program using a logistic regression model trained by RefSeq virus and prokaryote genomes. It takes k-mer features of a given contig as input and outputs the probability that the contig is a virus. Although these methods have been successfully used to uncover viral diversity from metagenomics^10^, both rely on similarity to known viral genomes. Therefore, as reference-based approaches, they cannot uncover viral lineages totally different from known viral genomes, if such lineages exist. Thus, we sought to develop a non-reference-based analysis pipeline to detect viral sequences from metagenomes; however, we were tasked with the question ‘What information could be used for this purpose’? The underlying biology of the clustered regularly interspaced short palindromic repeats (CRISPR) system, a prokaryotic form of adaptive immunological memory^11^, provides a potential source. After infection by viruses or horizontal transfer of plasmids, some archaeal and bacterial cells incorporate fragments of ‘non-self’ genetic materials in specialized genomic loci between CRISPR direct repeats (DRs). The incorporated sequences, called ‘spacers’, are identical in sequence to part of the previously infecting mobile genetic element. Thus, the genetic information encoded in CRISPR spacers can be inferred as likely viral, and distinguishable to the genetic material of the organism encoding CRISPR which is most often cellular, but potentially also viral^12,13^.

CRISPR spacers have been used to detect viral genomes^14–18^ and predict viral hosts^19,20^. They have previously been extracted from assembled bacterial genomes to assess CRISPR ‘dark matter’, finding that 80–90% of identifiable material matches known viral genomes^17,18^. In the current study, we extended this conceptual approach to enormous amount of unassembled short-read metagenomic data. CRISPR repeats are relatively easily identifiable, particularly compared to unknown viral sequences. This trait enables us to search immense metagenomic datasets for reads comprised in part as CRISPR DR sequences, and in other parts, unknown sequences inferred as CRISPR spacers. By analysing these reads and the contigs assembled from them, we successfully extracted 11,391 terminally redundant (TR) CRISPR-targeted sequences ranging from 894 to 292,414 bases. These sequences are expected to be complete or near-complete circular genomes which can be linked to their predicted hosts. The sequences we inferred de novo in this manner included 257 complete genomes of the crAssphage family^21,22^, 11 genomes larger than 200 kilobases (kb), 766 genomes of *Microviridae* species, 114 genomes of *Inoviridae* species and many novel, taxonomically unplaced genomes. Furthermore, by comparing the spacer-associated metagenomic CRISPR DR sequences to CRISPRs from reference bacterial and archaeal genomes, we predicted CRISPR targeting hosts for 69.7% of the discovered genomes.

## Results and Discussion

### Extraction of CRISPR-targeted sequences

For this study, we analysed human gut metagenomes as they serve as an ‘ecosystem’ with the most abundant metagenomic data available to our knowledge. We downloaded 11,817 human gut metagenome datasets equivalent to 50.7 Tb from the European Nucleotide Archive FTP server. FASTQ files were pre-processed and assembled to 180,068,349 contigs comprising 767.7 Gb of data (Supplementary Table 1). We discovered 11,223 unique CRISPR DRs from the assembled contigs which were used to extract CRISPR spacers from raw reads, resulting in 1,969,721 unique CRISPR spacers (Supplementary data). These spacers were then used as query to reveal candidate protospacers (i.e. contigs containing the spacer sequence not within a CRISPR locus). Spacers were mapped to CRISPR masked contigs with a 93% minimum sequence identity threshold. To increase specificity, we checked that the 5’ and 3’ adjacent sequences of spacer-mapped positions were not similar either to each other or the spacer-associated DR. A total of 164,590,387 candidate protospacer loci, attributed to 1,114,947 unique spacers (56.6% of all unique spacers) were revealed (Supplementary data), showing a substantially higher discovery rate compared with that of a previous study (∼7%^17^) which used National Center for Biotechnology Information (NCBI) nucleotide sequences for protospacer discovery. Spacers were clustered based on protospacer co-occurrence and used to extract contigs targeted by more than 30% of members of a spacer cluster. Finally, 764,883 gapless CRISPR-targeted sequences (15.9 Gb) were extracted (Supplementary Fig. 1 and 2). Of these, 11,391 unique sequences were identified as TR^23^, which are expected if originating as complete or near-complete circular mobile genetic elements. The size of CRISPR-targeted TR sequences ranged from 894 to 292,414 bases (Supplementary table 2 and Supplementary data).

We then investigated protein coding genes encoded in CRISPR-targeted sequences. To maximize recovery of novel genes, 3,099,178 protein coding genes were predicted from CRISPR-targeted sequences, including both TR (400,566 genes) and non-TR sequences (2,698,612 genes). Protein sequences were clustered based on a 30% sequence identity threshold, resulting in 83,796 clusters with more than two members and comprising 2,996,113 genes (96.7%). Each cluster was aligned and used to build a hidden Markov model (HMM), then used as query against UniRef50 and RefSeq viral proteins. We found that 12,283 HMMs (14.7%), comprising 891,121 genes (28.8%), were similar to 60,296 different RefSeq viral protein sequences (E-value < 10^-10). A total of 6,853 TR sequences (60%) encoded at least one of these genes, suggesting that a substantial portion of TR sequences are viral genomes.

### Length-based classification of CRISPR-targeted TR sequences

Evaluation of TR sequence length showed a multi-modal distribution with a distinct trough at 20 kb (Fig. 1a). For reference, we refer to the 8,837 TR sequences shorter than 20 kb as ‘small’ and the 2,554 TR sequences longer than 20 kb as ‘large’. This simple classification was previously used to infer capsid morphology^24,25^. Among large TR sequences, 1,509 sequences (59%) encode a set of tail, portal and terminase large-subunit genes (score > 30), whereas only 50 small TR sequences (0.6%) encoded this set of genes (Supplementary Fig. 3), indicating that the majority of large TR sequences are tailed phages, also known as *Caudovirales*. In addition, 766 small TR sequences, ranging in size from 4,574 to 12,988 bases (average 5,766.14 bases), encoded both *Microviridae* major capsid and replication initiator homologues^26^. 114 small TR sequences, ranging in size from 4,783 to 18,988 bases (average 6,715.6 bases), encoded both *Inoviridae* major coat and replication initiator homologues. We consider that this portion of small TR sequences are likely viruses with a non-tailed morphology^27,28^.

**Fig. 1.**
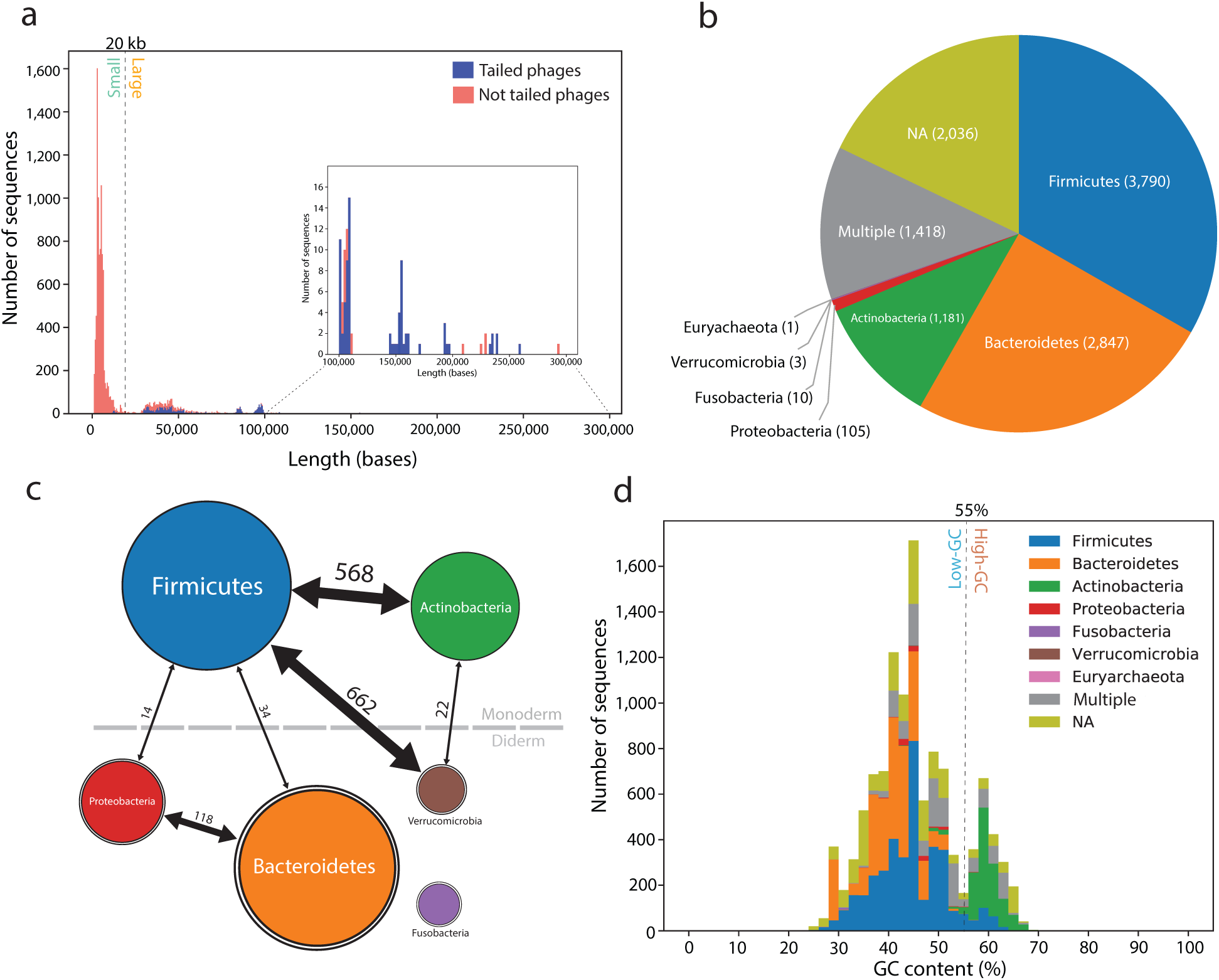
Properties of CRISPR-targeted TR sequences. **a**, Length distribution of TR sequences. We used portal, tail and terminase large subunit as marker genes of tailed phage; sequences encoding all three of these genes were categorized as tailed phages. The dotted line at 20 kb represents an arbitrary cut-off between small and large sequences. Sequences longer than 100 kb are shown in the inset. **b**, Predicted CRISPR targeting host composition of TR sequences. Hosts were predicted by mapping CRISPR DR sequences to the RefSeq database. Sequences containing ≥10 protospacer loci but less than 90% associated DR taxa exclusiveness were considered targeted by multiple phyla. If ≥10 protospacers could not be assigned to a taxon, the predicted targeting host is denoted as not available (NA). **c**, Heterogeneous distribution of TR sequences targeted by multiple phyla. Circle size approximately represents popularity of the respective host. Bidirectional arrows connect the top two host phyla according to host-assigned protospacer counts (i.e. protospacers most often associated with CRISPR DRs are assigned to these two phyla). Numbers on arrows are counts of the number of TR sequences targeted by the connected phyla. **d**, Predicted targeting host distribution by GC content. The dotted line indicates the low- and high-GC content boundary at 55%.

### Predicted CRISPR targeting hosts of TR sequences

As our approach uses CRISPR spacers to extract CRISPR-targeted sequences, we hypothesised that the relationship between virus and targeted host can be resolved for the majority of TR sequences. CRISPR DR sequences were searched on RefSeq genomes and taxonomically assigned. Based on counts of protospacers associated with taxonomically assigned DRs, 7,937 TR sequences (69.7%) were resolved to a targeting host at the phylum level (Fig. 1b and Supplementary Fig. 4), and 6,083 TR sequences (53.4%) were resolved to a targeting host at the order level (Supplementary Fig. 5 and Supplementary table 2). The most frequent host was *Firmicutes*, followed by *Bacteroidetes* and *Actinobacteria*. Notably, these are the most common bacteria in human intestine^29^. In addition, 1,418 TR sequences (12.5%) were predicted to be targeted by multiple phyla. Although some of these TR sequences were targeted by multiple monoderm or diderm phyla exclusively, there was an exceptional host ambiguity between *Firmicutes* and *Verrucomicrobia* which crosses the monoderm-diderm boundary (Fig. 1c). We did not use CRISPR DR sequences shared between cross-phyla species for targeting host prediction; targeting host ambiguity is unlikely because of ambiguous taxonomic assignment of DR sequences. To determine whether the targeting host predicted by this method corresponds with reported permissive hosts, we calculated GC content^30^ of the TR sequences (Fig. 1d). Among the 2,050 high-GC (GC% > 55%) TR sequences, 1,109 (54%) and 249 (12.1%) were predicted to be targeted by *Actinobacteria* and *Firmicutes* respectively, and the targeting host was undetermined for 222 TR sequences (10.8%). The large fraction of high-GC TR sequences predicted to be targeted by *Actinobacteria* likely indicate genomic adaption of parasitic genetic elements which infect and routinely become targeted by CRISPR systems of high-GC gram-positive bacteria.

Host range within a viral lineage was further assessed by phylogenetic analysis of TR sequences determined to represent *Microviridae* species. The predicted targeting hosts of putative *Microviridae* species from our study were *Bacteroides, Firmicutes* and *Proteobacteria*, indicating a wide host range of *Microviridae* species. Molecular phylogeny of the *Microviridae* major capsid protein clearly separates sequences based on their targeting host (Fig. 2). Interestingly, most known *E. coli*-infecting viral species (such as phiX174) and other *Proteobacteria*-infecting species (such as phi MH2K^31^) used as reference in this phylogeny split to two clades, and the clade containing the latter was within a clade of TR sequences targeted by *Firmicutes*. The nested phylogenetic structure of capsids encoded by TR sequences predicted to represent *Microviridae* species may indicate that host-switching, a critically important topic in virus evolution^32,33^, has occurred within the viral lineage.

**Fig. 2.**
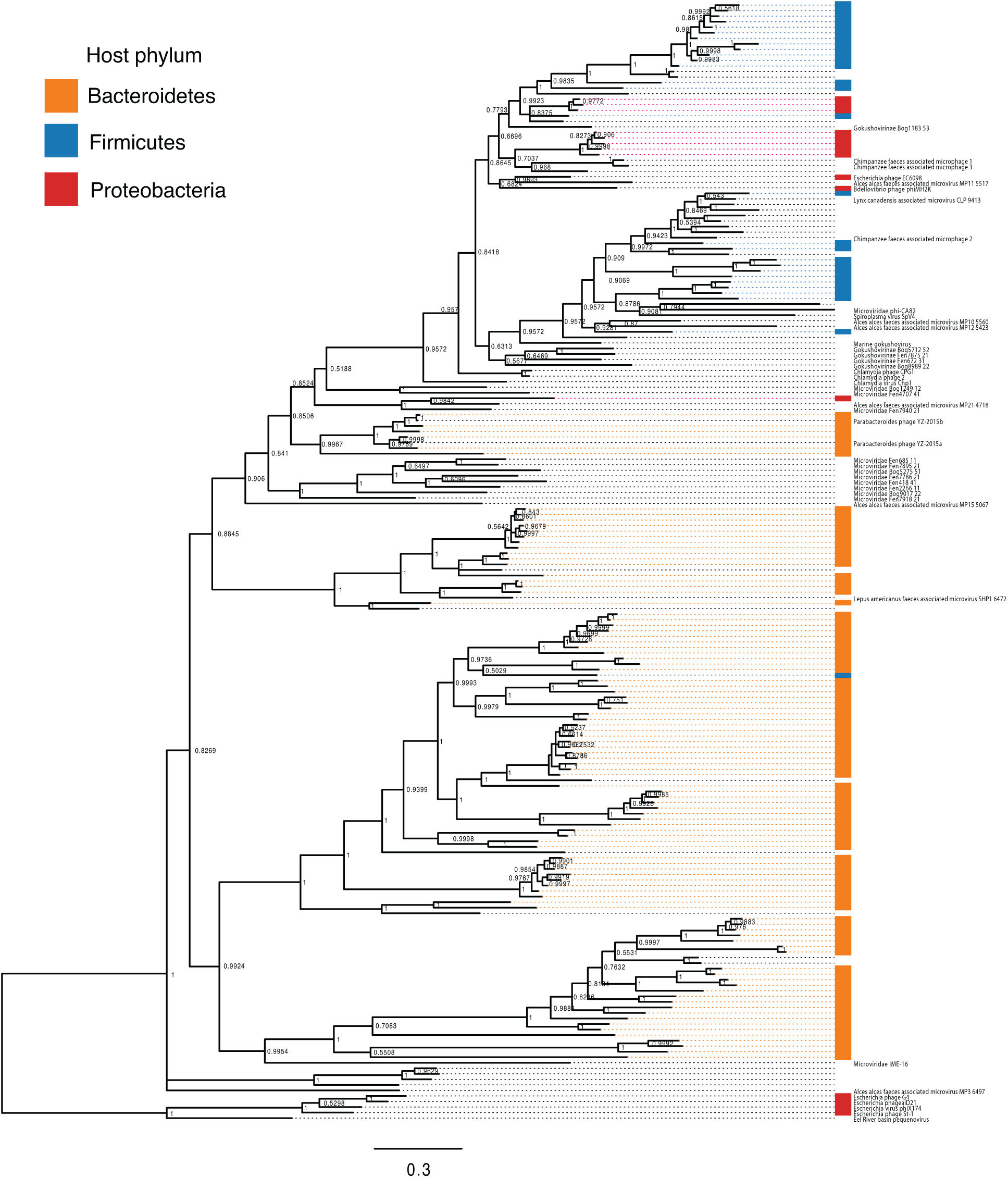
Bayesian phylogeny of *Microviridae* major capsid proteins. A total of 159 representative major capsid protein sequences from this study and 43 RefSeq major capsid protein sequences were used for analysis. Taxa without a name denote the *Microviridae* species from this study and taxa with text denote *Microviridae* species from RefSeq. Taxa were annotated based on predicted targeting hosts. The phi X174 clade was selected as the outgroup.

These results suggest that approximately 70% of discovered sequences are targeted by specific host phyla. 12.5% of TR sequences were targeted by multiple phyla; however, we are uncertain whether these elements actually infect multiple hosts at present, recently host-switched, or some genomes become CRISPR-targeted as a result of abortive infections^34^. Importantly, expanding this approach to detect viral genomes from different environments may be useful to infer the evolutionary history and genetic factors involved in host-switching.

### Comparison of CRISPR-targeted TR sequences to available viral and plasmid sequences

TR sequences identified in this study were compared to virus and plasmid genome databases, including RefSeq^35^ virus, RefSeq plasmid, IMG/VR^36,37^ and GVD^10^ (Fig. 3). IMG/VR is currently the largest database of uncultivated viral genomes. GVD is a database of viral genomes discovered from human gut metagenome datasets using VirSorter and VirFinder. Both of these databases rely on protein homology, which complements to our approach, to discover the viral genomes. Therefore, we consider these databases are appropriate to validate our results.

**Fig. 3.**
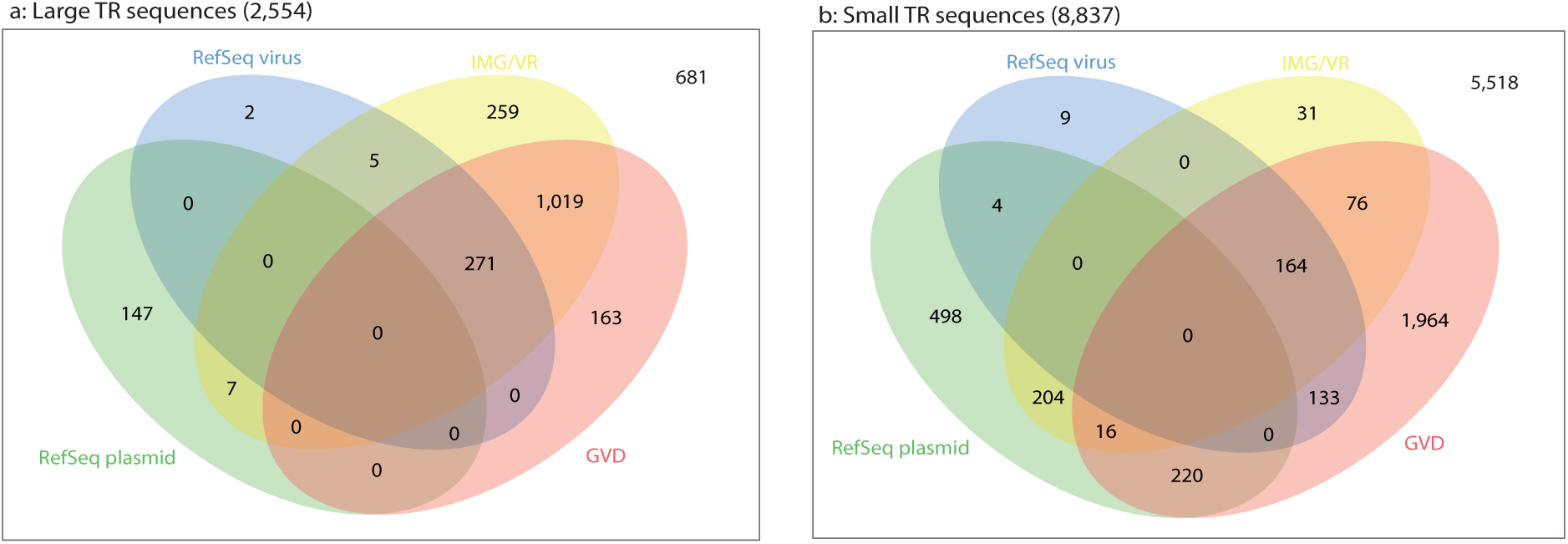
Venn diagrams of database comparisons for (a) large and (b) small TR sequences. Each TR sequence was compared to RefSeq virus, RefSeq plasmid, IMG/VR and GVD using BLASTN. Database hit minimum criteria was set to 85% sequence identity with 75% aligned fraction of query sequence to a unique subject sequence.

Among the 2,554 large TR sequences, we found that 1,726 TR sequences (67.6%) were represented in RefSeq virus, IMG/VR or GVD (Fig. 3a), using a threshold of 85% sequence identity with at least 75% aligned fraction of the query sequence to a unique subject sequence. These sequences included 257 crAssphage genomes, ranging in size from 92,182 to 100,327 bases (average 96,984.8 bases). Members of *Bacteroidales* were predicted as targeting hosts of crAssphages in this study, findings which are consistent with those hosts reported to propagate crAssphage in previous studies^21,22^. We also found that 154 large TR sequences (6%) were listed in RefSeq plasmid, of which 7 were also listed in the IMG/VR database. Based on our investigation, we determined that there is adequate classification between plasmids and viruses for large genomes. Notably, we discovered 11 TR sequences larger than 200 kb; 7 of which correspond to recently reported ‘huge phage’ genomes^13^ and 4 of these very large TR sequences were novel. Finally, we found that 681 large TR sequences (26.7%) were novel. In summary, we conclude that many of the large TR sequences are already represented in virus or plasmid databases, with the exception of those greater than 200 kb which were only recently reported.

In contrast to the large TR sequences, most of the small TR sequences were not represented in the databases searched (Fig. 3b). Among the 8,837 small TR sequences, 491 small TR sequences (5.6%) were listed in IMG/VR, whereas 2,573 small TR sequences (29.1%) were listed in GVD. Only 256 small TR sequences (2.9%) were listed in both IMG/VR and GVD. Finally, 5,518 small TR sequences (62.4%) were novel, indicating that the majority of small viral or plasmid sequences targeted by CRISPR are still unexplored, even in the intensively studied human gut metagenome. However, IMG/VR filters out genomes shorter than 5 kb to minimize the rate of false positive predictions^37^, which likely explains the substantially lower represented sequences in IMG/VR for small TR sequences. Moreover, among the 942 RefSeq plasmid-listed sequences, 444 small TR sequences were listed in RefSeq virus, IMG/VR and/or GVD, suggesting a possible misclassification between virus and plasmid within these databases. We investigated database representation of CRISPR-targeted TR sequences predicted to represent *Microviridae* and *Inoviridae* species. Among the 766 putative *Microviridae* genomes from this study, we found that 639 genomes (83.4%) were represented in at least one virus database, and none were listed among RefSeq plasmids. In contrast to *Microviridae*, among the 114 putative *Inoviridae* genomes from this study, only 2 were listed in the virus databases and 11 were listed in the RefSeq plasmid database. Next, we compared our putative *Inoviridae* genomes with recently reported *Inoviridae* genomes discovered using a machine learning approach^38^, and found that 43 genomes from our study were highly similar to the genomes from the previous study, supporting our prediction that these sequences are indeed *Inoviridae* genomes.

To determine whether our novel TR sequences are viruses, we searched sequences for capsid genes using a high-sensitivity homology detection method. We built a capsid HMM database using reference capsid proteins as baits which were then used to scan TR sequences. We identified 1,582 small TR sequences (17.9%) encoded capsid genes (score > 30), suggesting that at least this portion of TR sequences are viral genomes. However, capsid genes are unlikely to share a single common ancestor^39^, and it is unlikely that the reference sequences cover the entire sequence diversity of capsids. In addition, some viruses do not encode capsids^40,41^. Therefore, although this is a conservative estimate of the truly viral fraction of the taxonomically unclassifiable sequences reported here, we are unable to strongly conclude whether the remaining small TR sequences are viruses or not at this point. With DNA synthesis technology and predicted host information from this study, one might be able to experimentally investigate these potentially viral genomes.

### Gene content based hierarchical clustering of CRISPR-targeted TR sequences

To scrutinize CRISPR-targeted TR sequences in a genomic context, we hierarchically clustered TR sequences based on gene content. For this analysis, we selected the top 1,000 genes recurrently observed from large and small TR sequences. The clustering results for large TR sequences (Fig. 4a) showed that the majority of genomes, with a variety of gene contents, have already been listed in virus databases. This finding further supports that these databases contain at least representatives similar, at broad taxonomic levels, to the large viruses present in human gut. Specific gene contents were observed for crAssphages, which are exclusively targeted by *Bacteroidetes*. In addition, RefSeq plasmid hit sequences formed an exclusive cluster with conjugation-related genes. As the conjugation proteins included those involved in pili formation and intercellular DNA transfer proteins, these sequences are likely F plasmids. TR sequences predicted to be targeted by monoderm and diderm hosts clustered separately, indicating there is little gene flow between them. Except for the likely F plasmid sequences, most of these sequences encode capsids. Combined with the fact that most of the large TR sequences encoded tailed phage essential genes, this result further supports the conclusion that the majority of the large TR sequences are tailed phages.

**Fig. 4.**
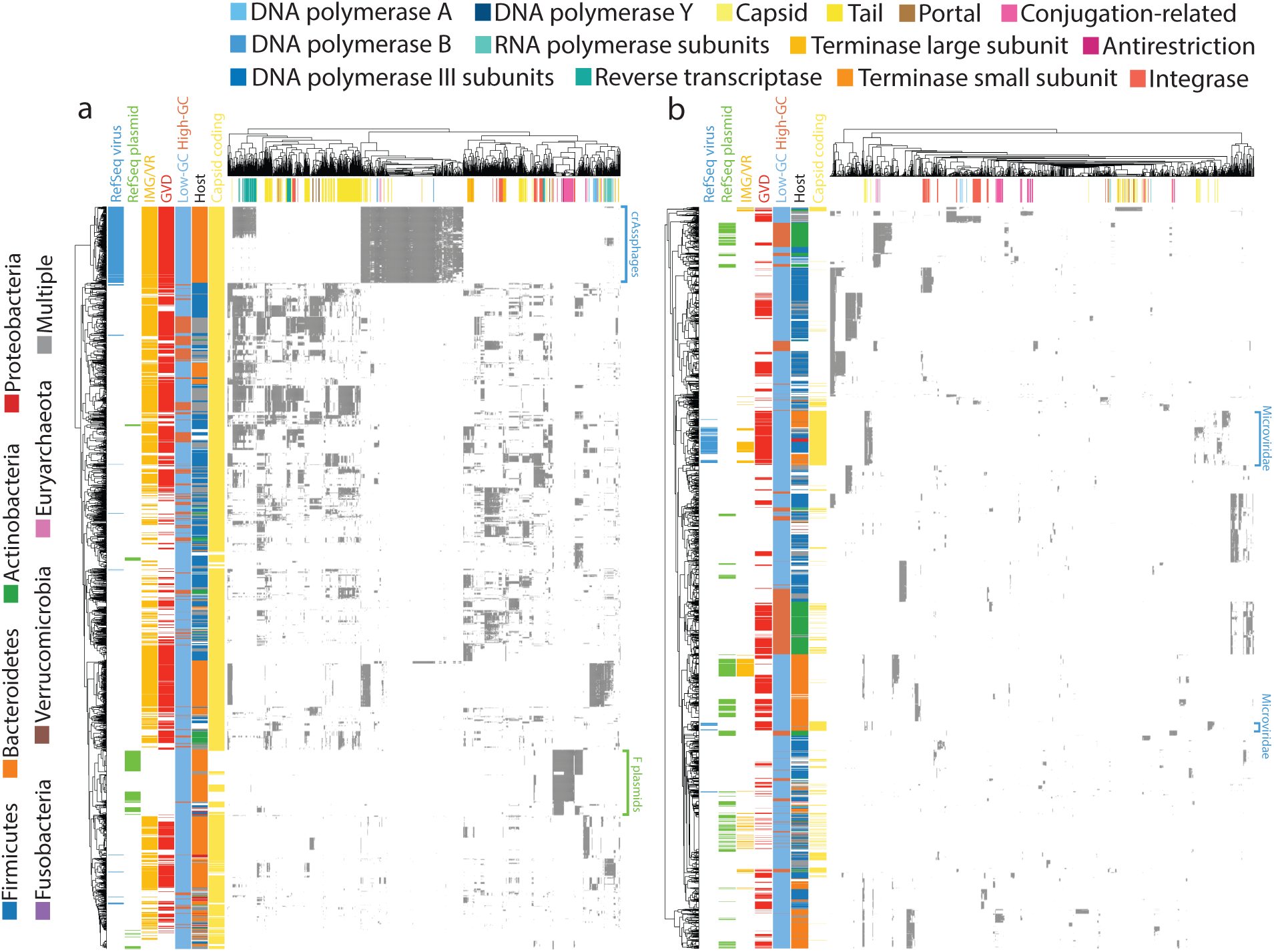
Hierarchical clustering of (a) large and (b) small TR sequences based on gene content. Heatmaps representing gene content of TR sequences, in which each row is a TR sequence and each column is a gene cluster. Gray areas in the heatmap indicate sequences encoding a gene that is homologous to the gene cluster. Note that one gene can be homologous to multiple gene clusters. Sequences are annotated by database containing similar sequences, GC content, host and detectable capsid genes. Gene clusters were annotated by searching corresponding HMMs to the UniRef50 database. Several notable RefSeq-listed clusters are denoted on the right side of the heatmaps.

In contrast to the large TR sequences, we found that the small TR sequences remained largely enigmatic when clustered by gene content. Few clusters were represented in IMG/VR, and although GVD covers a relatively broader range, representatives are still missing or sparse for some clusters. Several clusters were also listed in both plasmid and virus databases. As many clusters did not encode detectable capsid genes, both clusters representing *Microviridae* evince capsid genes. Another pattern shown by this analysis was that high-GC content TR sequences were frequently observed with *Actinobacteria* as the predicted targeting host.

### Diversity-generating retroelements of crAssphage lineage

Recently, one crAssphage genome encoding a diversity-generating retroelement (DGR) was reported^42^. DGRs are retroelements found in prokaryotes and viruses^43^. They minimally consist of an error-prone reverse transcriptase and a non-coding region called a template repeat. The transcript of a template repeat is reverse transcribed, and this complementary DNA (cDNA) product includes mutations from adenine to a random nucleotide (A-to-N). The cDNA is incorporated to a specific position of the target gene called the variable repeat; thus, this element effectively alters the coding sequence of the target gene which may benefit from rapid adaptation, such as expansion to new hosts and environments^44,45^. Among the 257 putative crAssphage genomes identified in our study, 98 genomes encoded DGRs including a reverse transcriptase gene with an associated non-coding region recognizable as a putative template repeat showing A-to-N (adenine-to-any base) mutations to the tail-collar coding gene. Using these abundant putative crAssphage sequences, we further investigated this locus to uncover the origin of this retroelement.

The position of the DGR locus was consistent among DGR-encoding crAssphages (Supplementary Fig. 6), suggesting that DGRs in crAssphages observed so far are orthologous and likely originate from a common ancestral DGR insertion. The reverse transcriptase gene and template repeat within this module showed high sequence conservation, whereas the variable repeat within the adjacent gene that encodes tail-collar fibre showed high diversity, indicating that this DGR was recently generating diversity (Fig. 5a). Notably, however, the tail-collar fibre of putative crAssphage sequences without DGR were similarly diverse. We used the protein sequence of DGR reverse transcriptase to query non-human-derived crAssphage-like genomes^46^, and only one crAssphage-like genome, which was derived by sequencing baboon faeces, encoded this element (Fig. 5a). Pair-wise alignment of template and variable repeats within the baboon crAssphage-like genome showed A-to-N substitutions that are likely facilitated by the DGR (Fig. 5b), which indicates the DGR in the baboon crAssphage-like genome is also functional. We then constructed a Bayesian phylogeny of crAssphage genomes (Fig. 5c). We found that the distribution of reverse transcriptase-encoding genomes (shown in red text in Fig. 5c) did not form a cluster, a finding which is most consistent with multiple losses of this retroelement, and potentially explaining the high diversity of the tail collar fibre even in assembled sequences that do not currently contain a DGR. A deeper-rooted molecular phylogeny with distant crAssphages-like genomes would be required to uncover more details about the origin of this retroelement.

**Fig. 5.**
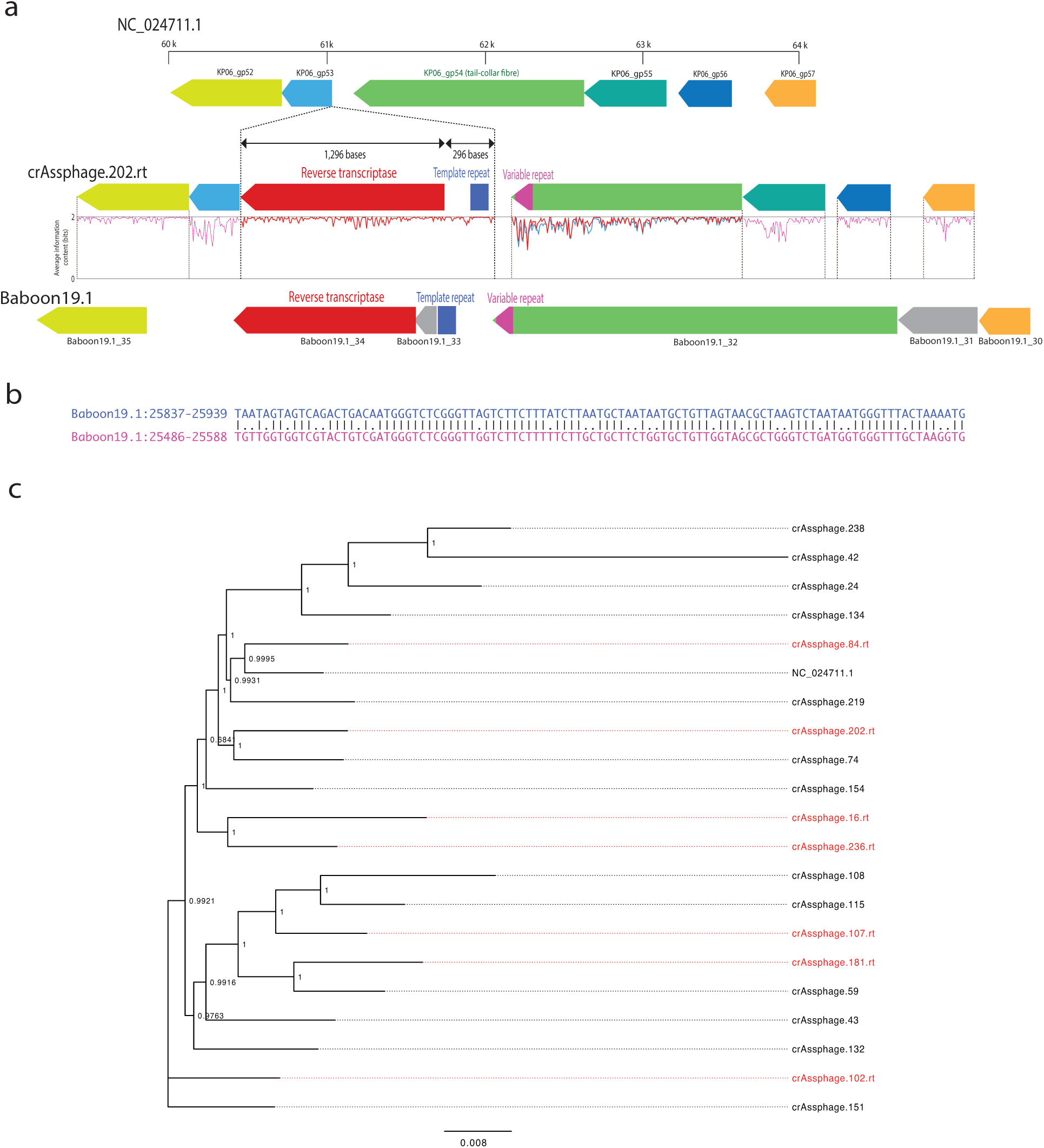
DGR locus and phylogeny of crAssphage core genes. **a**, Comparison of the DGR locus between a reference crAssphage genome (NC_024711.1), a representative from this study (crAssphage.202.rt) and a baboon-derived crAssphages (Baboon19.1). The retroelement-insertion region is enclosed by a dotted line. Sequence conservation of DGR and its target gene (tail collar fibre) were calculated by aligning and building HMMs. Sequence conservation of the DGR target gene was calculated separately based on whether the genome encodes DGR (red) or not (blue). Information content was averaged over seven base-window sizes. **b**, Pairwise sequence alignment of template (top) and variable (bottom) repeats from Baboon19.1. **c**, Unrooted Bayesian phylogeny of crAssphage core genes. 55 core genes from 20 representative crAssphages genomes and one reference genome were used for analysis. Those crAssphages with DGR appear in red.

### Remnant CRISPR spacers and overall contribution of CRISPR-targeted sequences to identified spacers

Viruses and other mobile genetic elements can escape from CRISPR targeting by acquiring mutations to protospacer loci^47,48^. While the corresponding spacers are thus no longer effective, they can remain in the host genome. To investigate these potential ‘remnant’ CRISPR spacers, we mapped all unique CRISPR spacers to TR sequences as well as scrambled sequences with various sequence identity thresholds (Fig. 6). The scrambled sequences were used to monitor false positive matches arising by chance (see Methods). Based on the observation of incremental false positive matches of spacers to scrambled sequences, 84% is a sequence identity threshold at which some putative remnant spacers can be mapped, but very few false-positive matches. At an 84% sequence identity threshold, 269,808 spacers and 126,616 spacers were mapped to large and small TR sequences, respectively. Taken together, 20.13% of all unique spacers (396,424 spacers) were mapped to TR sequences. Compared to the stringent identity threshold initially applied (93%), 91.91% more spacers were mapped to small TR sequences with relaxed threshold. By comparison, only 42.68% more spacers were mapped to large TR sequences with relaxed threshold. These results suggest that a substantial fraction of CRISPR spacers imperfectly matches to protospacers within circular mobile genetic elements in human gut. This potentially reflects an ‘escape mutation’ phenomenon. By percentage, small TR sequences can be explained in this manner than large TR sequences.

**Fig. 6.**
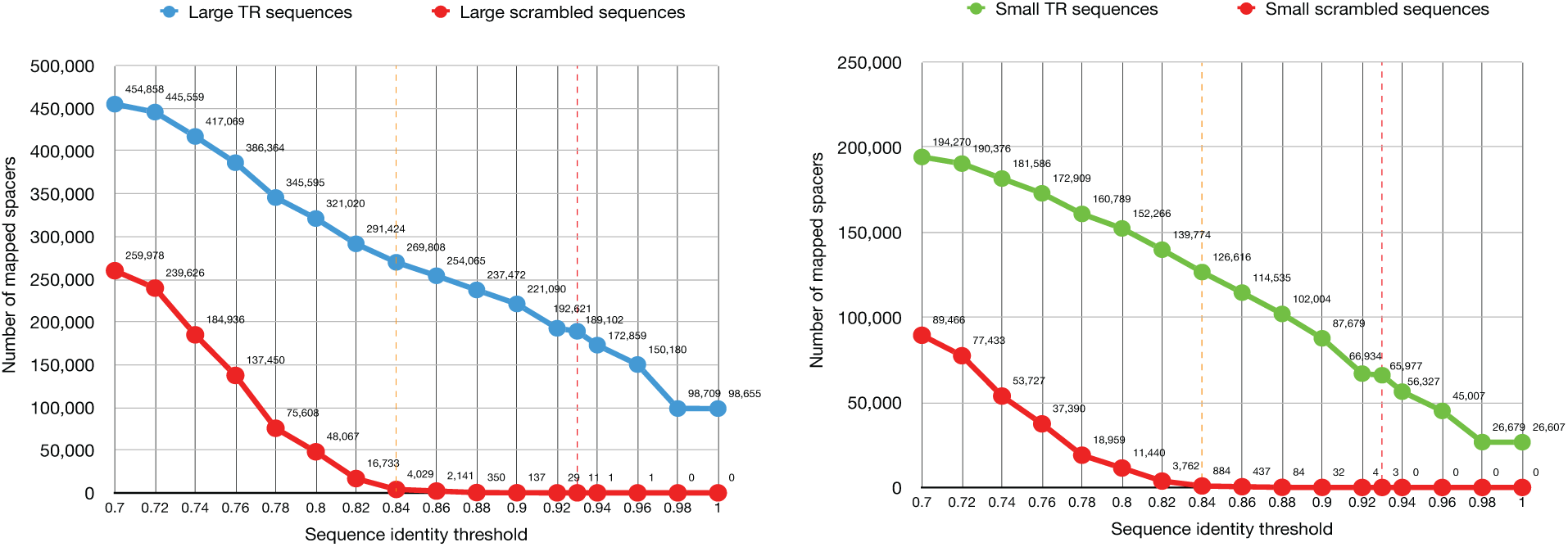
Number of mapped spacers according to various sequence identity thresholds. All unique CRISPR spacers were mapped to large TR sequences, small TR sequences and scrambled sequences. The initially applied and relaxed sequence identity thresholds are denoted as red and orange colored dashed lines respectively. The spacer mapping process is identical to the protospacer discovery process (see Methods).

While we focused TR sequences in this study because of high confidence of the genomic completeness, spacers could be derived from incompletely reconstructed or non-circular genomes. To estimate the contribution of discovered CRISPR-targeted sequences to identified CRISPR spacers, all unique spacers were mapped to all representative CRISPR-targeted sequences with 84% sequence identity threshold. Under these conditions, 971,224 spacers (49.31% of all unique spacers) were mapped.

Overall, spacer mapping with looser criterion suggesting that at least about one-fifth of discovered CRISPR spacers originated from TR sequences or their recognizable evolutionary predecessors, whereas about half of CRISPR spacers originated from our discovered CRISPR-targeted sequences including both TR and non-TR.

## Conclusion

In the current study, we demonstrated that CRISPR spacers can be used as a tool to detect viral genomes and other mobile genetic elements from metagenome sequences. Using spacers to confidently infer which sequences are targeted by CRISPR, we substantially expanded the diversity of mobile genetic elements identifiable from the human gut metagenome, which has already been a topic of intense investigation for virus discovery. Comparing the sequences predicted by this approach against viral databases showed that our protocol effectively detects viral genomes without requiring similarity to any known viral sequence. Although the majority of large (>20 kb) genomes were predicted as tailed phages with high confidence based on sequence homology, we found that the majority of small (<20 kb) genomes remain unclassified because of a lack of similar genomes in annotated databases. We showed that the source of nearly half of the CRISPR spacers encoded by residents of the human gut remains unknown, suggesting that additional protospacer reservoirs, whether extinct or simply unsampled, are still to be characterized. Applying this conceptual advance to additional metagenomic datasets will increase the breadth of the lens through which we can study the diversity of Earth’s virome.

## Materials and Methods

### Materials

Sequencing data were selected based on NCBI metadata. Filtering parameters for the query were as follows: layout = PAIRED, platform = ILLUMINA, selection = RANDOM, strategy = WGS, source = METAGENOME, NCBI Taxonomy = 408170 (human gut metagenome) and minimum library size = 1 Gb. If a sample contained multiple runs, we selected the run with the most bases for simplicity of the analysis pipeline and to avoid possible bias to protospacer counts from nearly identical metagenomes.

### Database versions and download dates

RefSeq Release 98 was downloaded on 10 January 2020 and IMG/VR Release Jan. 2018 was downloaded on 21 October 2019. GVD was downloaded on 11 March 2020 and UniRef50 was downloaded on 16 December 2019.

### Metagenome assembly

All downloaded paired FASTQ files were pre-processed based on guidance in BBTools^49^ (version 38.73). Adapters, phi X and human sequences were removed using BBDuk and BBMap. Sequencing errors were corrected using Tadpole. Each pre-processed pair of FASTQ files was assembled using SPAdes^50^ (version 3.12) with -meta option. Contigs smaller than 1 kb were discarded.

### Detection of CRISPR and spacer extraction

Assembled contigs were scanned with CRISPRDetect^51^ (version 2.2) to extract CRISPR DRs which were deduplicated using CD-HIT-EST^52^ (version 4.7) and used to mask the raw reads using BBDuk. We extracted CRISPR spacers from the raw reads to maximize spacer capture from the library. Sequences between the masked regions within the raw reads were considered CRISPR spacers which were extracted by a simple Python program (available in our source code repository) and then deduplicated.

### Detection of protospacer loci

All DRs were mapped to contigs using BBMap with a 93% minimum sequence identity. DR mapped positions and their flanking 60 bases were masked as CRISPR loci. Next, identified spacers were mapped to all CRISPR masked contigs with a 93% minimum sequence identity. Rather than excluding all contigs with CRISPR loci, we exclusively masked CRISPR loci to identify viruses that encode CRISPR systems which are themselves targeted by other CRISPR systems. To increase specificity, we aligned the 5’ and 3’ adjacent regions of spacer-mapped positions. These adjacent sequences were also aligned to the DR sequence associated with the mapped spacer. We discarded those loci in which any alignment score divided by length was higher than 0.5, using the alignment parameters: match = 1, mismatch = −1, gap = −1 and gap extension = −1. The remaining positions were considered authentic protospacer loci.

### Co-occurrence-based spacer clustering

We clustered spacers in two steps (Supplementary Fig. 2). First, we clustered protospacer loci located within 50 kb of another protospacer locus. We then further clustered spacers based on the co-occurrence of protospacers represented as a graph. In this graph, protospacers are nodes and the edges represent co-occurrence of connected protospacers. The weights of edges were observed counts of co-occurrence of the connected protospacers defined in the previous clustering. Graph communities were detected using Markov Clustering algorithm^53^ (options: -I 4 -pi 0.4; version 14-137). Clusters with a size lower than 10 and a global clustering coefficient lower than 0.5 were discarded. Finally, 12,749 clusters comprising 591,189 spacers were derived.

### Extraction of CRISPR-targeted sequences

Contiguous regions of contigs targeted by more than 30% of members of a spacer cluster were marked as a bed file using BEDTools^54^. To join the fragmented clusters, adjacent regions within 1 kb were concatenated. Marked regions were extracted, and sequences containing assembly gap were discarded. Finally, both ends of each extracted sequence were compared to identify TR sequences using a python program utilizing the Biopython^55^ package (available in our source code repository).

### Deduplication of CRISPR-targeted sequences

TR sequences were clustered using PSI-CD-HIT (options: -c 0.95 -aS 0.95 -aL 0.95 -G 1 -g 1 -prog blastn -circle 1). The remaining CRISPR-targeted sequences were clustered twice using linclust^56^ (options: --cluster-mode 2 --cov-mode 1 -c 0.9 --min-seq-id 0.95), then clustered again using PSI-CD-HIT (options: -c 0.9 -aS 0.95 -G 1 - g 1 -prog blastn -circle 1).

### Gene prediction and annotation of CRISPR-targeted sequence

Protein-coding genes were predicted from both TR and non-TR sequences using Prodigal (version 2.6.3) with the -p meta option. To recover truncated genes from TR sequences, each TR sequence was concatenated in silico and unique predicted genes were selected. Predicted protein sequences with partial frags were discarded. The remaining protein sequences were clustered based on a 30% sequence identity threshold and aligned using mmseqs^57^ (version e1a1c1226ef22ac3d0da8e8f71adb8fd2388a249). HMMs were constructed from each aligned sequence using hmmbuild from HMMER^58^ (version 3.2.1). Constructed HMMs were then used as query to search against UniRef50 and RefSeq viral proteins (E value < 10e-10) using hmmsearch.

### Capsid HMM database construction

We selected capsids and coat proteins from RefSeq viral protein sequences. Sequences containing ‘capsid’ and ‘coat’ in the description were selected and then manually inspected to remove sequences unlikely to actually be capsid, such as containing the term ‘associate’ or phrase ‘assembly chaperone’. A total of 5,069 sequences were clustered based on a 30% sequence identity, resulting in 1,324 representative sequences. Each representative sequence was used as bait and underwent five jackhmmer iterations on UniRef50 database. HMMs from the final iteration were used to build the capsid HMM database.

### Targeting host prediction

DRs were mapped to RefSeq bacteria and archaea genomes using BBMap. A locus with more than three consecutive DR hits within 100 bases was considered an authentic CRISPR locus associated with the mapped DR. DRs mapped to multiple taxa in a given taxonomic level were not taxonomically assigned for that level. DRs assigned to taxa were used to predict the targeting host. We counted the protospacers linked to taxonomically assigned DRs within a TR sequence. If the count of a given taxon was ≥10 and higher than 90% exclusiveness, we considered that the corresponding taxon is a targeting host of a given contig. Host prediction was done for each taxonomic level: species, genus, family, order, class, phylum and domain.

### Gene content-based hierarchical clustering of TR sequences

TR sequences were scanned by HMMs derived from the clustering results of the predicted protein sequences using both TR and non-TR sequences. The scanned result was represented by a binary matrix (score > 60). We selected the top 1,000 genes recurrently observed within TR sequences. The matrix was hierarchically clustered with ComplexHeatmap^59^ (version 2.5.3), and then annotated with database hits, host, GC content, capsids and predicted gene functions. Clustering was separately performed for large and small TR sequences.

### Phylogenetic analysis of *Microviridae* major capsid proteins

Representative *Microviridae* major capsid protein sequences were selected by clustering all capsids based on an 85% sequence identity threshold throughout the entire length of the protein. Representative and reference protein sequences were aligned using MAFFT^60^ (version 7.310) and then trimmed using trimAl^61^ (version 1.4). Aligned sequences were used for Bayesian phylogenetic analysis using MrBayes^62^ (version 3.2.7). A mixed substitution model with a uniform prior that converged to Blosum62 (posterior probability = 1.000) was selected. All other priors were set to default. Two chains of Markov chain Monte Carlo (MCMC) with identical priors run over ten million generations and sampled every 500 generations. The standard deviation of split frequencies approached close to zero (0.007837) over the run.

### Phylogenetic analysis of crAssphage core genes

55 core crAssphage genes which were encoded in all crAssphage genomes from our study and the reference crAssphage genome were selected. Representative crAssphage genomes were selected by clustering RNA polymerase coding sequences based on a 97.5% sequence identity threshold. Each gene was aligned using MAFFT and then trimmed using trimAl. Aligned core genes were then concatenated and used for Bayesian phylogenetic analysis using MrBayes. A GTR + I + Γ substitution model was selected. Two chains of MCMC with identical priors were run over ten million generations and sampled every 500 generations. The standard deviation of split frequencies approached close to zero (0.000499) over the run.

### Generation of scrambled sequences

Scrambled sequences are random sequences with identical lengths to the TR sequences. The sequences are generated based on sampled nucleotide frequencies from the TR sequences using Biopython package (available in our source code repository).

## Code availability

The source codes used in this study are available at https://github.com/ryota-sugimoto/virome_scripts

## Data availability

Data set including discovered CRISPR spacers, direct repeats, protospacers, co-occurrence-based spacer clustering result, predicted protein sequences, built HMMs, database comparison results, phylogenetic analysis results and CRISPR-targeted TR sequences are available in our google drive. (URL: https://drive.google.com/drive/folders/1QwpG3TilpBFwmOD5_qi2NESWA_XC0_g0?usp=sharingMD5:d0f2ebe8dd20de4033a68735015943d4)

## Acknowledgements

The majority of analysis has been done on the supercomputer systems in National Institute of Genetics (NIG) and Okinawa Institute of Science and Technology Graduate University. R.S. is supported by funds of Database Center for Life Science, Research Organization of Information and Systems, and MEXT KAKENHI Grant Number 20K21405. R.S. thanks to NIG Research Administrator room for scientific discussions.

## Author contributions

R.S. designed the studies and performed most of the analyses. L.N. assisted in sequence alignment and phylogenetic analysis. P.N.T. assisted in metagenome assembly. J.I. gave scientific input in the data analyses and interpretation of the results. N.F.P. gave scientific input and helped write the manuscript. H.M., K.K and H.N. provided scientific advice on the entire study. I.I. designed the study and wrote the manuscript together with R.S.

