## Supplementary figures for "De novo virus inference and host prediction from metagenome using CRISPR spacers"

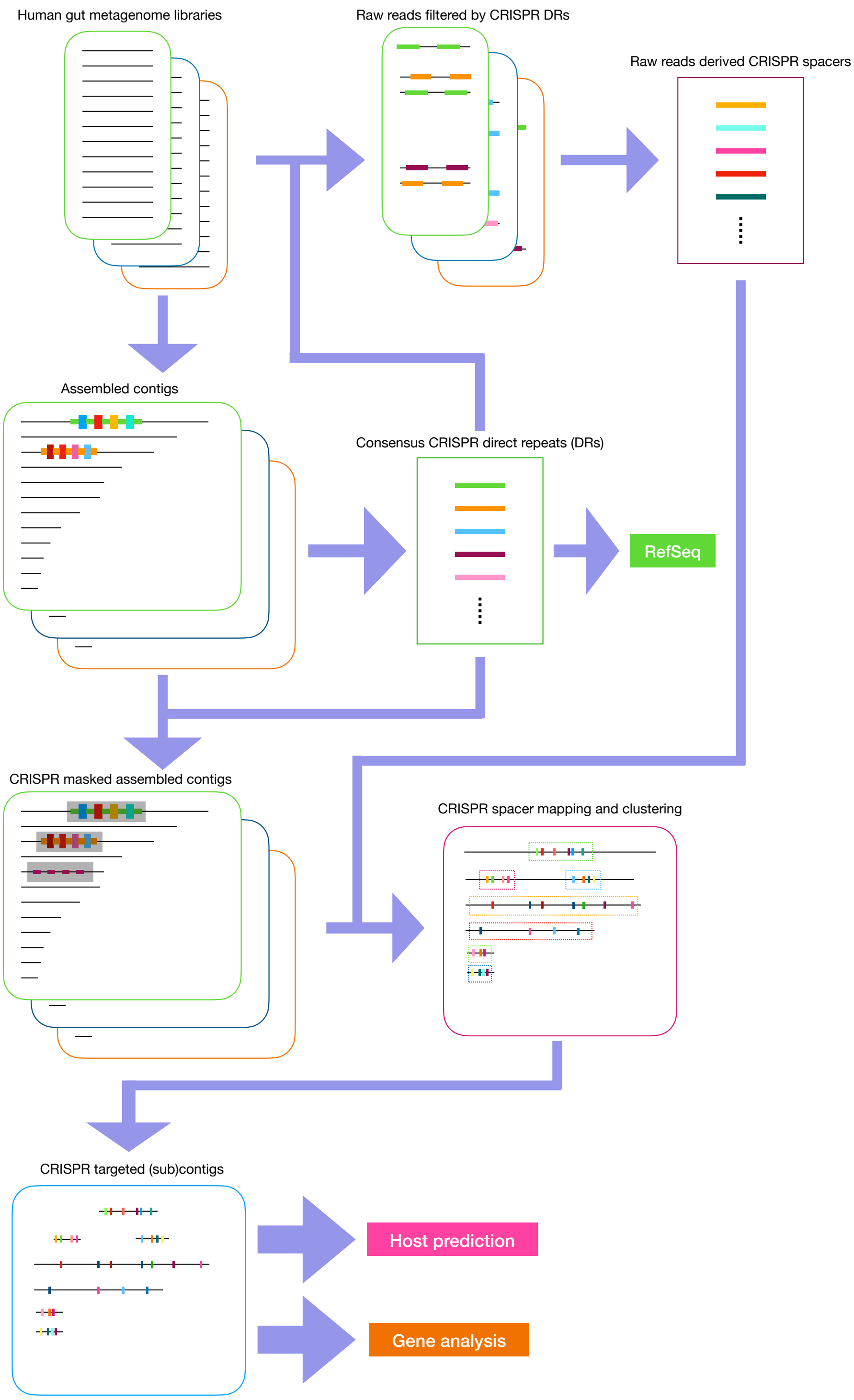


**Supplementary Fig. 1 | Basic workflow for viral genome detection.** Human gut metagenome libraries were pre-processed to remove adapters, phi X and human sequences. After correcting sequencing errors, libraries were assembled. Clustered regularly interspaced short palindromic repeats (CRISPR) loci were discovered from the assembled contigs. Consensus direct repeats (DRs) from discovered CRISPR loci were used to extract spacers, mask the CRISPR loci and predict the host. All unique CRISPR spacers were mapped to contigs to discover the protospacer loci. Spacers were clustered based on co-occurrence of the associated protospacers. Sequences targeted by more than 30% of members of a spacer cluster were extracted and used for further analysis.

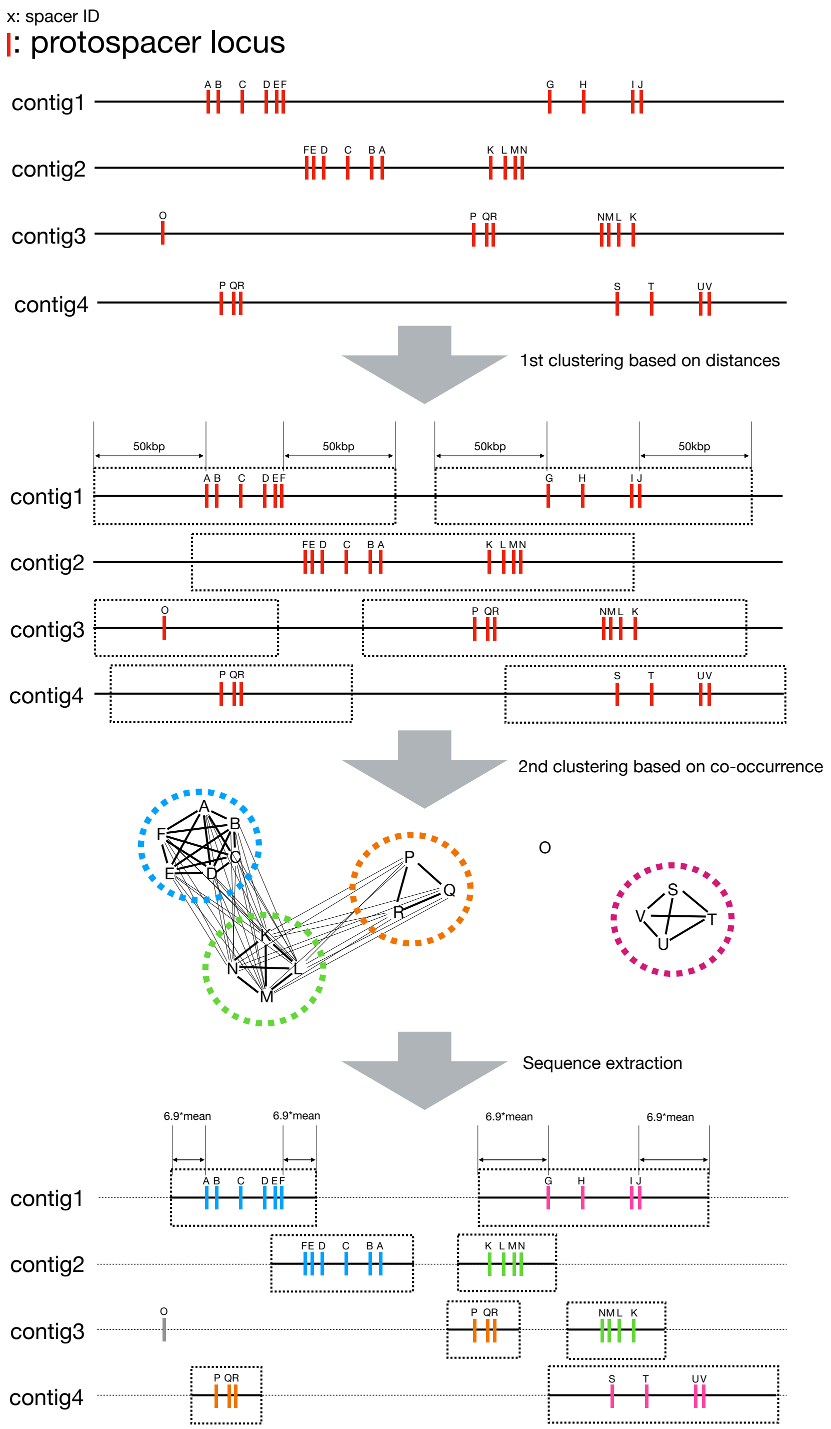


**Supplementary Fig. 2 | Spacer clustering based on co-occurrence of protospacers.** Initially, protospacer loci were clustered based on the distance between them. Within initial clusters, co-occurrences of protospacers were counted and used to construct an undirected graph. The nodes (spacers) in the undirected graph were further clustered using Markov clustering algorithm. Mean distances between adjacent protospacer loci within clusters were calculated and used to extract CRISPR-targeted sequences. The length and number of protospacers shown here are conceptual and not based on observed data.

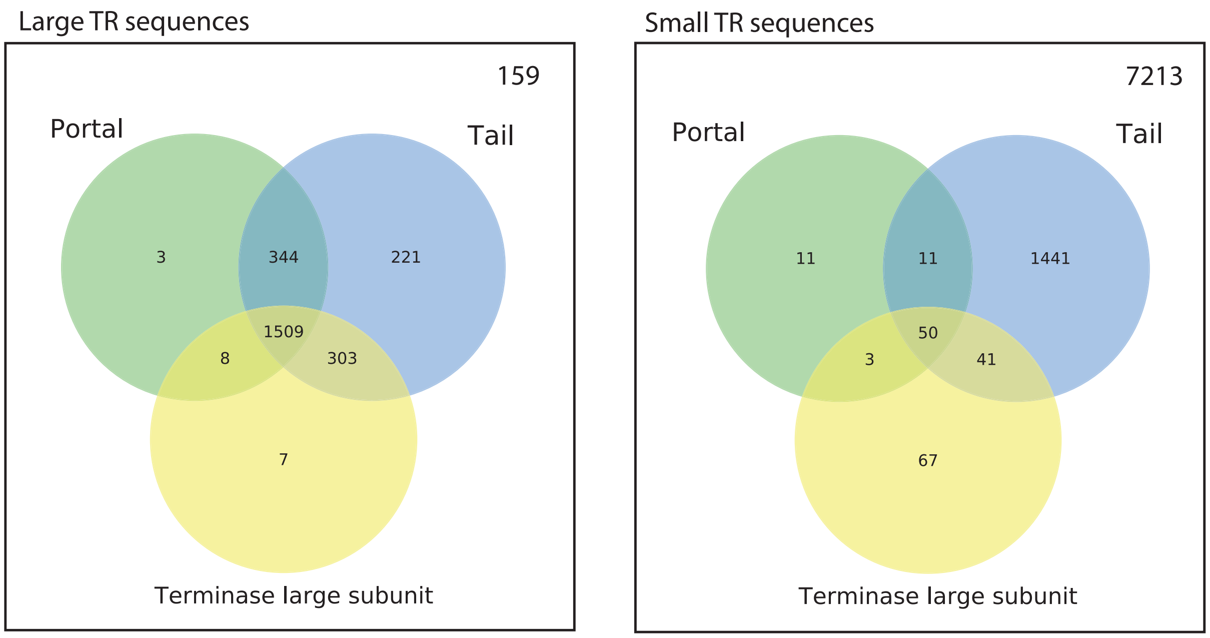


**Supplementary Fig. 3 | Venn diagrams of tailed phage marker genes.**

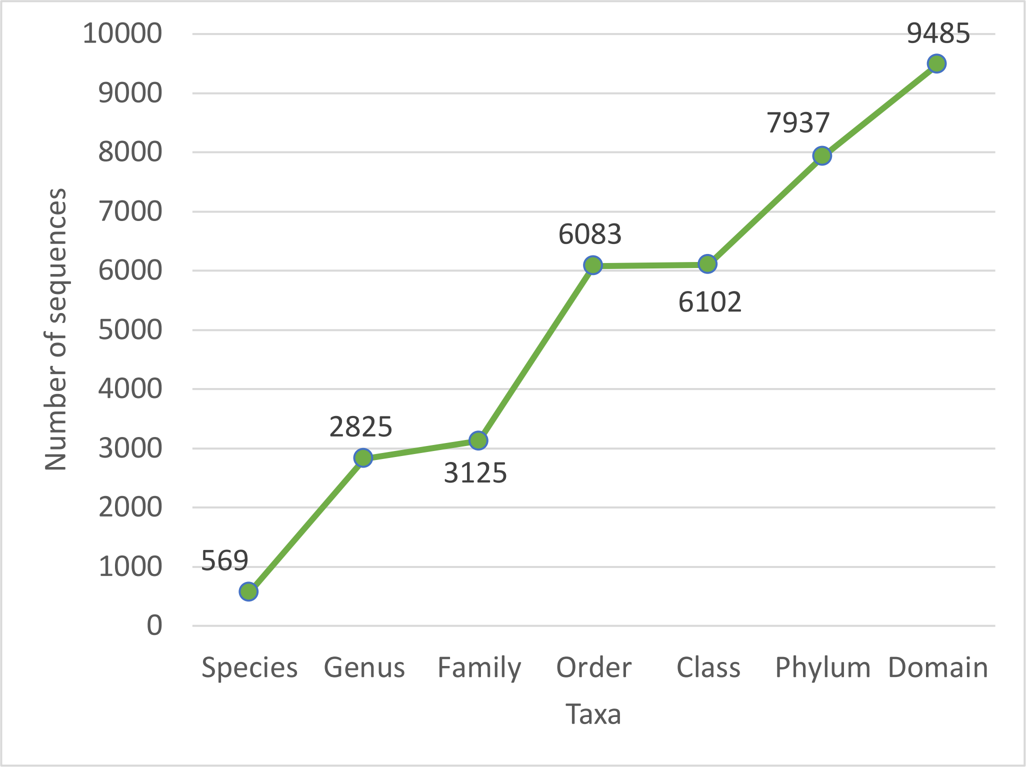


**Supplementary Fig. 4 | Number of sequences with a predicted targeting host according to each taxonomic level.**

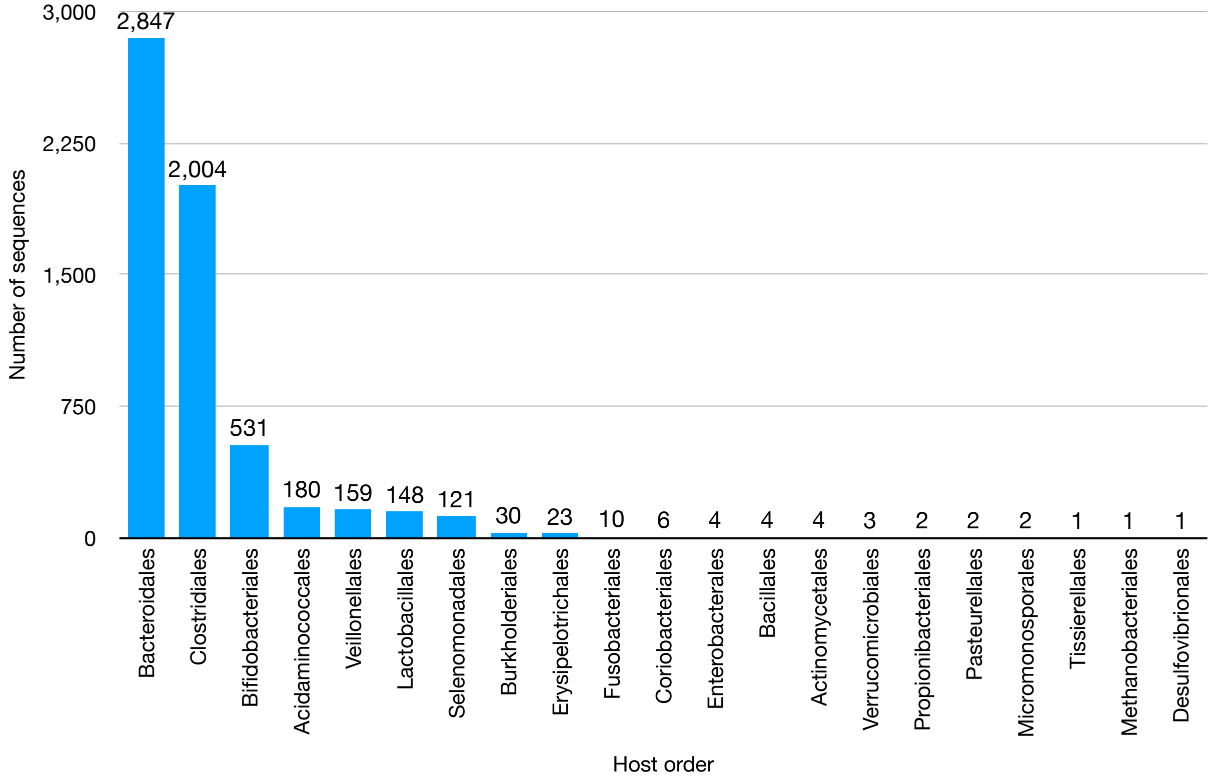


**Supplementary Fig. 5 | Number of sequences with a predicted CRISPR targeting host at the taxonomic level of order.**

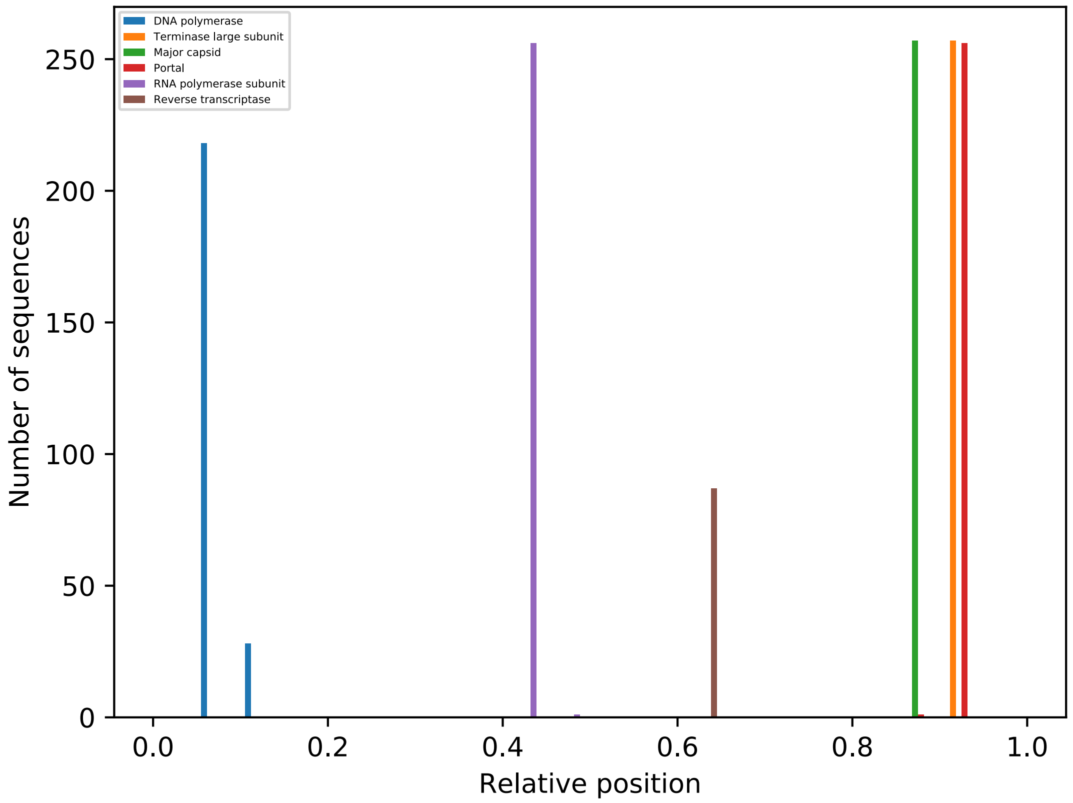


**Supplementary Fig. 6 | Relative positions of crAssphage essential genes and reverse transcriptase gene.** DNA polymerase, terminase large subunit, portal and RNA polymerase are selected as the essential genes. To define an origin of the circular genomes, the sequences from our study were aligned to the reference sequence NC_024711.1, then circulated in silico to phase with the reference. Positions were normalized by contig lengths, and gene loci were derived by searching contigs using the protein sequences as query.
